# Development of a copper-graphene nanocomposite based transparent coating with antiviral activity against influenza virus

**DOI:** 10.1101/2020.09.02.279737

**Authors:** Indrani Das Jana, Partha Kumbhakar, Saptarshi Banerjee, Chinmayee Chowde Gowda, Nandita Kedia, Saikat Kumar Kuila, Sushanta Banerjee, Narayan Chandra Das, Amit Kumar Das, Indranil Manna, Chandra Sekhar Tiwary, Arindam Mondal

## Abstract

Respiratory infections by RNA viruses are one of the major burdens upon global health and economy. Viruses like influenza or coronaviruses can be transmitted through respiratory droplets or contaminated surfaces. An effective antiviral coating can decrease the viability of the virus particles in the outside environment significantly, hence reducing their transmission rate. In this work, we have screened a series of nanoparticles and their composites for antiviral activity using Nano Luciferase based highly sensitive influenza A reporter virus. Using this screening system, we have identified copper-graphene (Cu-Gr) nanocomposite shows strong antiviral activity. Extensive material and biological characterization of the nanocomposite suggested a unique metal oxide embedded graphene sheet architecture that can inactivate the virion particles only within 30 minutes of pre-incubation and subsequently interferes with the entry of these virion particles into the host cell. This ultimately results in reduced viral gene expression, replication and production of progeny virus particles, slowing down the overall pace of progression of infection. Using PVA as a capping agent, we have been able to generate a Cu-Gr nanocomposite based highly transparent coating that retains its original antiviral activity in the solid form.

## Introduction

The emergence of novel virus strains and the associated outbreaks are becoming a significant threat to mankind (Koven 2020). The currently ongoing pandemic, caused by the Severe Acute Respiratory Syndrome-Coronavirus 2 (SARS-CoV-2), has brought the majority of the world to a grinding halt, severely impacting health & economy across the nations (Letko, Marzi, and Munster 2020). So far, the COVID-19 pandemic has claimed 720,000 lives resulting from 19.4 million infections globally. This reminds us of the great 1918 Spanish flu pandemic by the influenza virus, which resulted in 500 million infections with 50 million deaths (Johnson and Mueller 2002) (Patterson and Pyle 2019) (Taubenberger and Morens 2006) (Landrigan et al. 2018). Till date, there have been four worldwide pandemics caused by influenza viruses in the last century (1918, 1957, 1968, 2009) (World Health Organization 2018) (Saunders-Hastings and Krewski 2016), while Covid19 is the first pandemic caused by the Coronavirus (Y. Wu et al. 2020). Other than pandemics, influenza A and B viruses cause seasonal outbreaks (290,000 to 650,000 deaths worldwide: CDC FLUVIEW] and different coronaviruses cause mild flu-like symptoms to severe respiratory infections (SARS (De Wit et al. 2016) and MERS (Lin et al. 2019) coronavirus epidemics during 2002 and 2012 respectively) (Fehr and Perlman 2015). Clearly, recurring respiratory infections caused by these viruses are becoming one of the global human health problem.

Both influenza and coronaviruses cause respiratory infections, which can be transmitted from infected to healthy individuals through respiratory droplets, aerosols or contacts. These respiratory pathogens are known for their ability to persist on inanimate surfaces for days and even up to months, depending upon weather conditions (Vasickova et al. 2010). As a result, touching contaminated surfaces in public places is a potential route of viral transmission. In fact, the inanimate surfaces have been identified as a major cause of infections, especially in institutions where individuals are in contact with patients or contaminated fomites. Thus, the development of low cost and easily scalable antiviral coating materials, which could be widely applied to various surfaces in order to inactivate the virus particles in the environment, may serve as an effective way to reduce the chance of infection and hence to lower the overall speed of transmission.

Different metal oxides, including Cu and Ag have been explored for their biocidal activity in soluble as well as in insoluble forms (Minoshima et al. 2016). Copper and silver nanoparticles have remarkable properties like high electrical and thermal conductivity (Barani et al. 2020; D. Deng et al. 2013), superior catalytic nature (Gawande et al. 2016), anti-fungal (Cioffi et al. 2005) and bacteriostatic activities (Ruparelia et al. 2008) to name a few. Solid-state cuprous oxide and silver nitrate have been shown to inactivate virus particles through interfering with the activity of the surface antigen, hemagglutinin (HA), thereby blocking the attachment of the virus to the host cell receptors (Fehr and Perlman 2015) (Sunada, Minoshima, and Hashimoto 2012). Cuprous oxide nanoparticles had also been shown to inhibit the attachment and entry stages of Hepatitis C Virus infection, hence suggesting a generic mechanism for the copper-based nanoparticles for their antiviral activities (Hang et al. 2015). This may also explain the lesser viability of infectious SARS-CoV-2 particles on copper surfaces (8 h) when compared to stainless steel and plastic surfaces (72 h) (Neeltje van Doremalen et al. 2020). Biological activity of the Cu or Ag nanoparticles largely depends upon their size, shape, stability and capping agent. Due to the high reactivity of these nanoparticles, they can undergo rapid agglomeration leading to the drastic reduction of their activity (Ma et al. 2011). One way to overcome this instability is to form a composite with other organic or inorganic compounds which stabilizes the nanoparticles by altering their surface architecture (Perdikaki et al. 2018).

Graphene is composed of a single atom thick sheet of sp2 hybridized carbon atoms that forms a honeycomb lattice (Mohammed et al. 2020; Palmieri and Papi 2020). This structure of graphene is responsible for its large surface area, excellent electrical conductivity, strong mechanical strength and unique physicochemical properties (Edwards and Coleman 2013; Eigler and Hirsch 2014; Novoselov et al. 2012). It is used extensively in the field of nano-medicine due to easy surface functionality and controlled selectivity (Park et al. 2020). In recent times, the two dimensional sheet of graphene has caught much attention due to its antimicrobial and antiviral activity (Georgakilas et al. 2012). Differentially functionalized graphene oxide sheets can warp and encapsulate microorganisms, thereby severely restricting their interaction with host cells (Perdikaki et al. 2018) (Vinothini and Rajan 2017) (Karahan et al. 2018). For example, sulfate functionalized reduced graphene oxide has been shown to interact with the positively charged surface proteins of different orthopoxvirus strains (Ziem et al. 2016) (Ye et al. 2015). In this work, we have performed an extensive investigation of the potential antiviral activity of copper (Cu) nanoparticles, silver (Ag) nanoparticles, graphene (Gr) and their hybrid versions (nanocomposite materials) Ag-graphene, Cu-graphene and Ag-Cu-graphene against respiratory viruses using influenza A virus as a model system. Our data shows that prior incubation with the colloidal Cu-Gr nanocomposite can impose a strong reduction in viral infectivity, which gets manifested in the reduced viral entry, gene expression and subsequent production of progeny virions. Extensive material characterization using UV-Visible absorption and Raman spectroscopy along with X-Ray Diffraction (XRD) reveals a unique architecture of copper oxide decorated two-dimensional graphene sheets. The shape, size and distribution of the hybrid nanoparticles were also studied using Scanning Electron Microscopy (SEM) and Energy-dispersive X-ray spectroscopy (EDAX). Finally, we have developed for the first time a polyvinyl alcohol (PVA) based copper-graphene nanocomposite coating, which is completely transparent and shows strong antiviral activity in the solid phase.

## Results

### Synthesis and preliminary characterization of the nanoparticles and nanocomposites

We have synthesized various nanoparticles and their hybrids using a simple and cost-effective chemical method as discussed in the experimental section. The synthesized materials are Cu nanoparticles, Ag nanoparticles, graphene (Gr) and their hybrid versions Ag-graphene, Cu-graphene and Ag-Cu-graphene (Figure 1 A, B). Visible absorption spectra of each of these variants confirms the presence of respective components either alone or in combination with their composite partners, as shown in Figure 1C and D. The peak at 418 nm confirms the formation of Ag colloidal nanoparticles (Jin et al. 2005). From literature, it is attributed to Surface Plasmon Resonance with spherical nature (Hu et al. 2013). From the UV-vis spectra of the composite Ag-Gr, Plasmon absorption band at around 410 nm, indicates the formation of a hybrid structure. Incorporation of Ag nanoparticles on graphene sheets had led to a blue shift of the surface plasmon resonance, a characteristic similar to previous reports (Kim et al. 2018).

**Figure 1.**
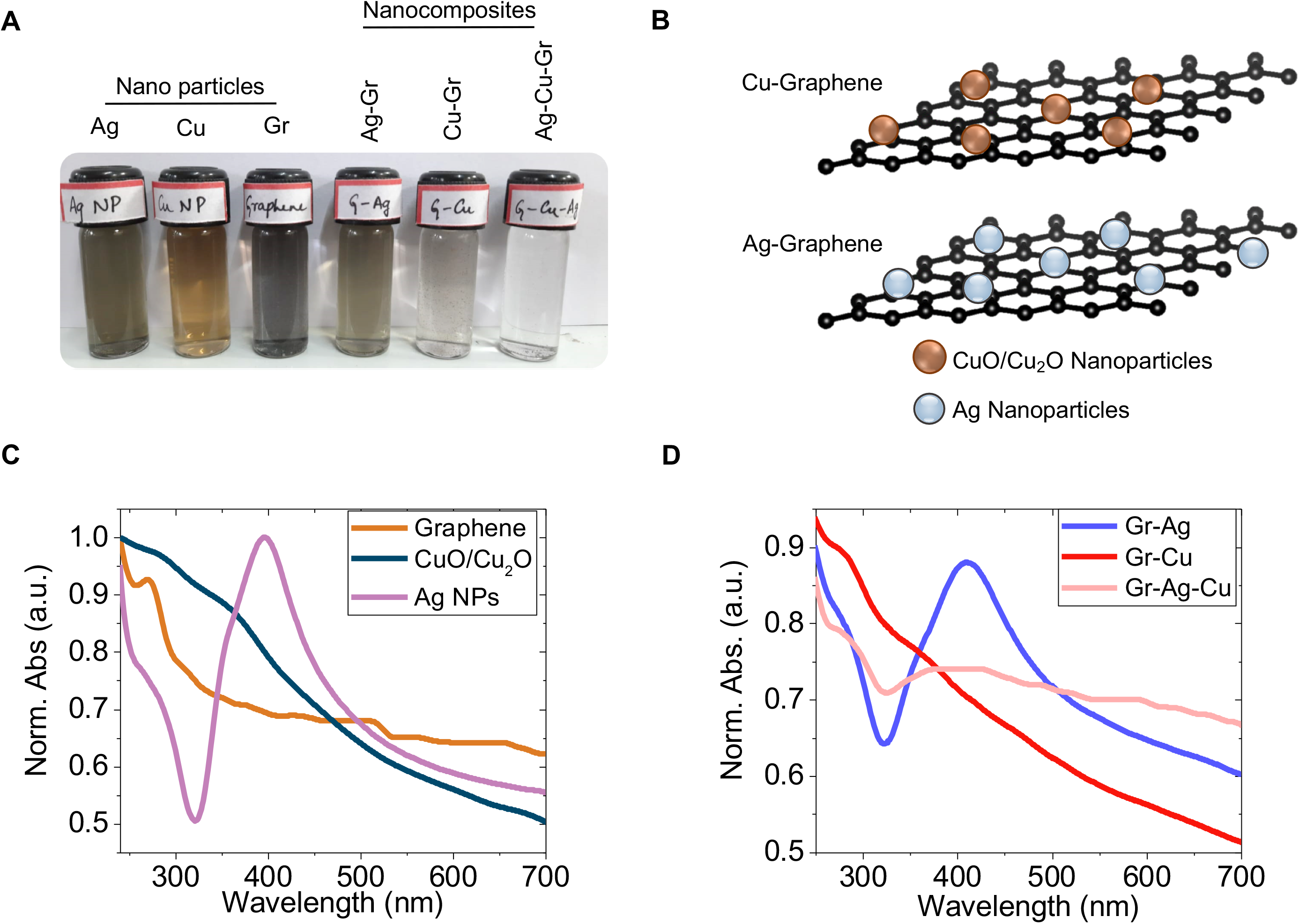
Different nanoparticles and their nanocomposites. (A) Digital photographs of all the synthesized nanoparticle and nanocomposite samples. (B) Schematic representation of the Cu and Ag nanoparticles embedded two dimensional graphene sheets. (C-D) UV-Vis absorption spectra of synthesized nanoparticles and their composites.

Furthermore, Cu nanoparticles, due to exposure of air at room temperature, goes through surface oxidation, which is the reason that samples contain oxygen in +1 and +2 states forming CuO or Cu_2_O (Yao et al. 2005). Composite including both Ag and Cu nanoparticles were also analyzed from the absorption spectrum. A broad graphene peak was observed at 280 nm in the composite. Both Ag and Cu nanoparticle plasmon peaks were observed and confirmed the presence of the colloidal nanoparticles on graphene sheets (Darabdhara et al. 2017). Particularly, Figure 1D shows the absorption spectra of the composites and it confirms the formation of graphene and CuO/Cu_2_O nanoparticles. An absorption peak at 262 nm is due to π→π* transition in C=C bond of graphene. Another absorption band at~350-450 nm is attributed to the intrinsic band to band transition of the CuO and Cu_2_O (Zhang et al. 2020). A blue shift was observed in the peak position compared to bulk CuO; this might be due to the quantum confinement effects exhibited by the particle when the size varies from bulk to nano. These UV-Vis peaks were well matched with CuO and Cu_2_O absorption profiles from previous studies (Chan et al. 2007). A broadened plasmon resonance peak for Cu-Gr nanocomposites is due to irregular shapes and sizes of the particles (Zhang et al. 2020).

### Screening of nanoparticles and their hybrids for antiviral activity

To test the antiviral activity of synthesized nanomaterials and their composites we have used a bioluminescent reporter variant of the influenza A virus, strain A/H1N1/WSN/1933, that has been previously reported by Tran et al. (Tran et al. 2013). This virus has a Nano-Luciferase (NLuc) gene fused to the carboxy-terminal of the viral PA gene, interspaced by the “self-cleaving” 2A peptide encoding sequence from porcine teschovirus. The Nano-Luc-influenza A reporter virus, as a part of its gene expression, synthesizes the PA-2A-NLuc polypeptide, which gets self-cleaved to produce Nano-luciferase. Subsequently, the luciferase activity could be measured as a quantitative estimate of viral gene expression and hence progression of virus replication cycle inside the cells. To test whether the Nano-luciferase activity could actually serve as a proxy to virus replication, we have infected MDCK cells with different amounts of input virus and viral replication/gene expression was monitored using Nano-Glo assay (Promega). As shown in Figure 3A, there is a linear relationship between multiplicity of infection (0.01-0.1) and luciferase light unit measurements (R^2^= 0.9294), where an increase in one log in the input virus amount leads to about 50% increase in the luciferase activity or vice versa, measured at 8 hours of post-infection. This data suggests that the Nano-luciferase influenza A reporter virus could serve as an excellent tool to study the antiviral activity of various nanoparticles or their nanocomposites used in this study.

In order to test the antiviral activity, we have standardized a “Nano-Luc reporter assay” described in Figure 3B. Briefly, Nano-luciferase influenza A reporter viruses were pre-incubated with the 5uM colloidal suspensions of each of the nanoparticles/ composites or with the vehicle control for 30 minutes at room temperature and subsequently used to infect MDCK cells at an MOI of 0.1. Luciferase activity was measured at 8 hours of post infection and plotted as a relative percentage of the vehicle control set (Figure 3C). Prior treatment of the virus stock solution with Cu-Gr composite showed 64% reduction in viral gene expression, while prior treatment with Ag-Gr resulted in 20% reduction. Treatment with other materials shows no significant decrease in luciferase activity. From the correlation of input virus units and the corresponding luciferase activity, as shown in Figure 3A, it can be inferred that prior treatment with Cu-Gr solution resulted in more than 10-fold reductions in the infectious virus population that has been used to infect the MDCK cells. In this context, it should be noted that none of the materials showed substantial cytotoxicity upon Madin-Darby Canine Kidney cells (MDCK) within the concentration range of 0.5 uM - 5.0 uM as evaluated using MTT assay (Figure 2). Hence, the reduction in Nano-luciferase activity as a result of prior exposure to Cu-Gr should be attributed exclusively to the reduction of the infectivity of the Nano-luciferase reporter virus. Henceforth, we focused upon the extensive characterization of the Cu-Gr nanocomposite.

**Figure 2.**
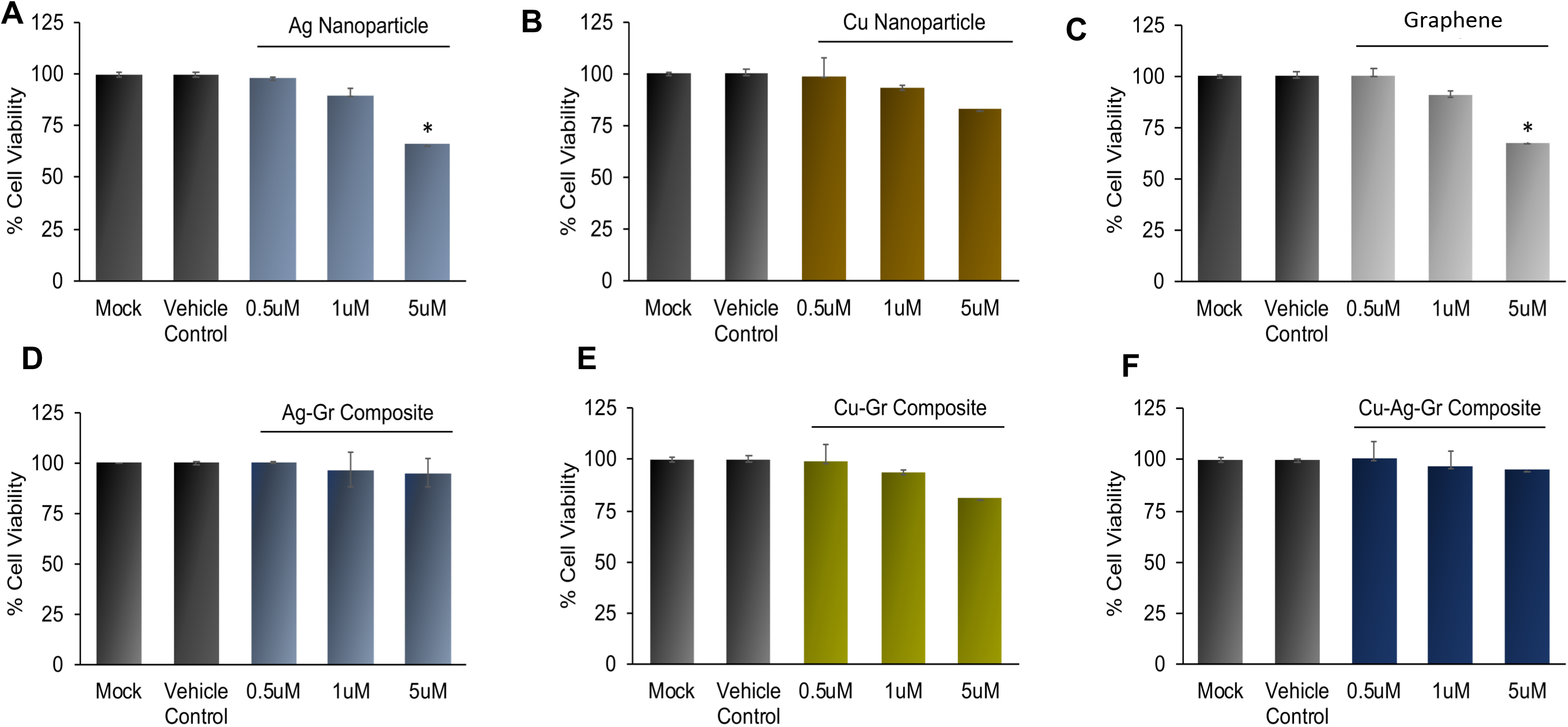
Nanoparticles and nanocomposites shows minimal cytotoxicity. MDCK cells were either treated with vehicle control or different concentrations (0.5, 1, 5 μM) of the Ag, Cu and Gr nanoparticles or Ag-Gr, Cu-Gr and Ag-Cu-Gr nanocomposites for 24 h and their cytotoxicity were determined by MTT assay. Cellular cytotoxicity was determined in triplicate and each experiment was repeated three times. Data are presented as means ± standard deviations (SD) (n=3).

**Figure 3.**
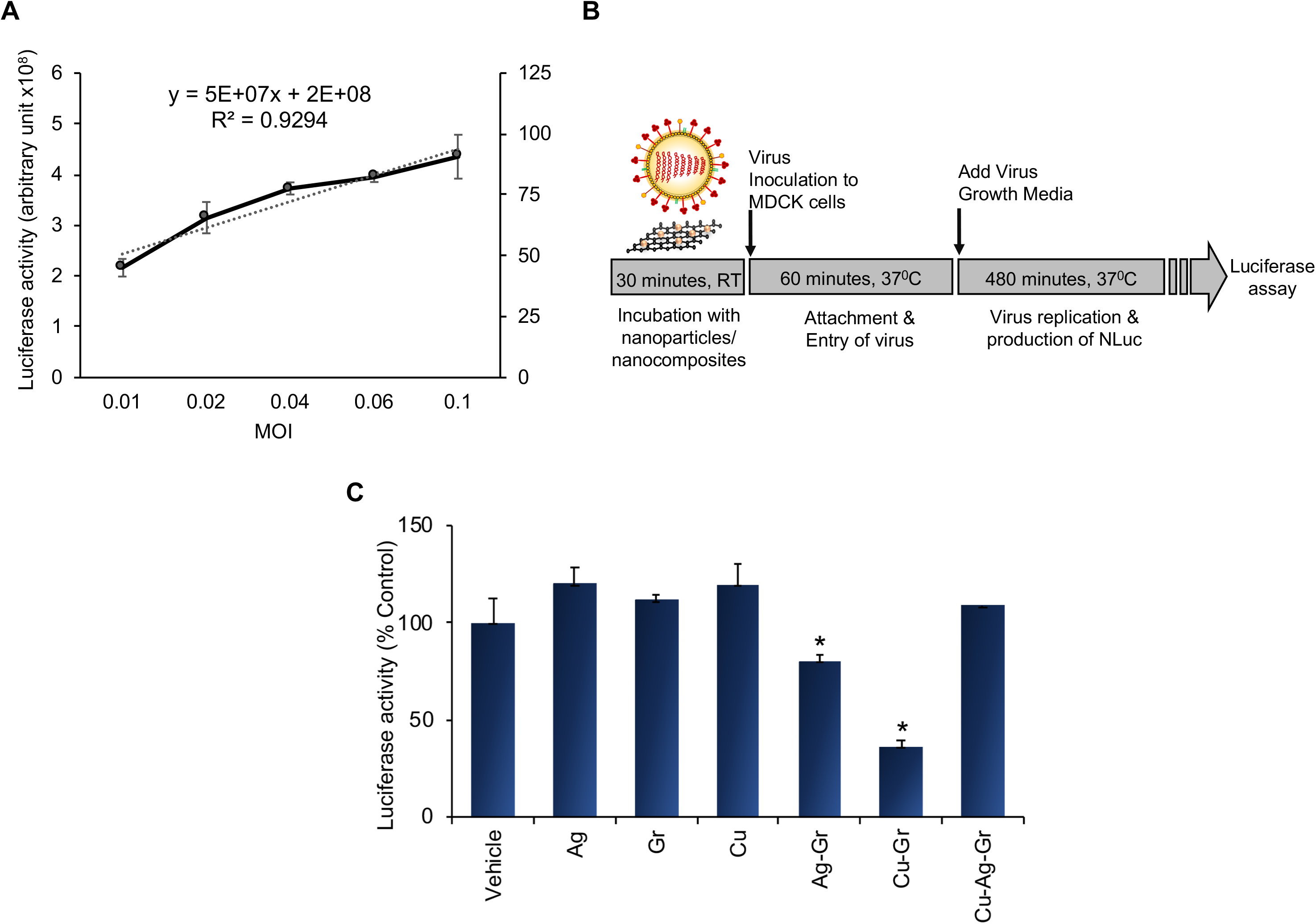
Screening of different the nanoparticles/ nanocomposites for antiviral activity using Nano-Luciferase influenza A reporter. (A) MDCK cells were infected with different amounts (0.01-0.1) of Nano-Luciferase Influenza A reporter virus and the reporter activity was measured using Nano glo reporter assay (Promega). To demonstrate the relationship between the MOI and luciferase activity (arbitrary unit) was plotted as a function of MOI. (B) To test the antiviral activity, the Influenza A reporter virus was pretreated with the 5uM colloidal suspensions of each of the materials for 30 minutes at RT and followed by infection of MDCK cells at MOI of 0.1as diagrammed. (C) Luciferase activity of the nanoparticle/ nanocomposite treated sets were measured at 8 hpi and plotted as a relative percentage of the vehicle treated set. For all experiments, data are mean of n=3± standard deviation (*, P< 0.05; n=3 ± sd).

### Material characterization of Cu-Gr nanocomposite

We have extensively characterized the structural parameters of the synthesized Cu-Gr nanocomposites by optical measurements. Figure 4 A depicts Raman spectra of synthesized Cu-Gr nanocomposite samples at excitation of 532 nm in the range of 200 cm^−1^ to 3000 cm^−1^. With Raman spectroscopy, we are able to distinguish both pristine graphene and copper peaks. The presence of D peak (1361 cm^−1^) and G peak (1527 cm^−1^) confirms the existence of graphene in the samples synthesized. Generally, D peak originates from defects in the hexagonal sp^2^ carbon system while the G peak arises due to the stretching vibration of sp^2^ carbon pairs in both rings and chains (Ferrari et al. 2006). Except, D and G peak, the 2D peak arises at ~2700 cm^−1^. The 2D peak originates due to transverse optical (TO) phonons around the K point and is activated by triple resonance Raman scattering (TRRS) (J. Bin Wu et al. 2018). In the measured Raman spectra (Figure 4B), three peaks are (280 cm^−1^, 350 cm^−1^ and 654 cm^−1^) observed to confirm the formation of oxide of Cu and originate due to the first order phonon scattering. The peaks are assigned to A_g_ and 2B_g_ peaks of copper oxide (Y. Deng et al. 2016). The graphene sheets are also seen in the optical images, as shown in the inset of Figure 4 A. Figure 4B shows the XRD patterns of graphene, CuO, and Cu_2_O nanoparticles, which confirm the crystalline phase of composites samples (standard JCPDS file number 35–0505, 80–1917). Interestingly, the intensity of the Cu_2_O is more intense compared to that of the CuO peaks, suggesting a higher abundance of Cu_2_O on the surface of the graphene sheets. No diffraction peaks corresponding to impurities are observed in the patterns.

**Figure 4.**
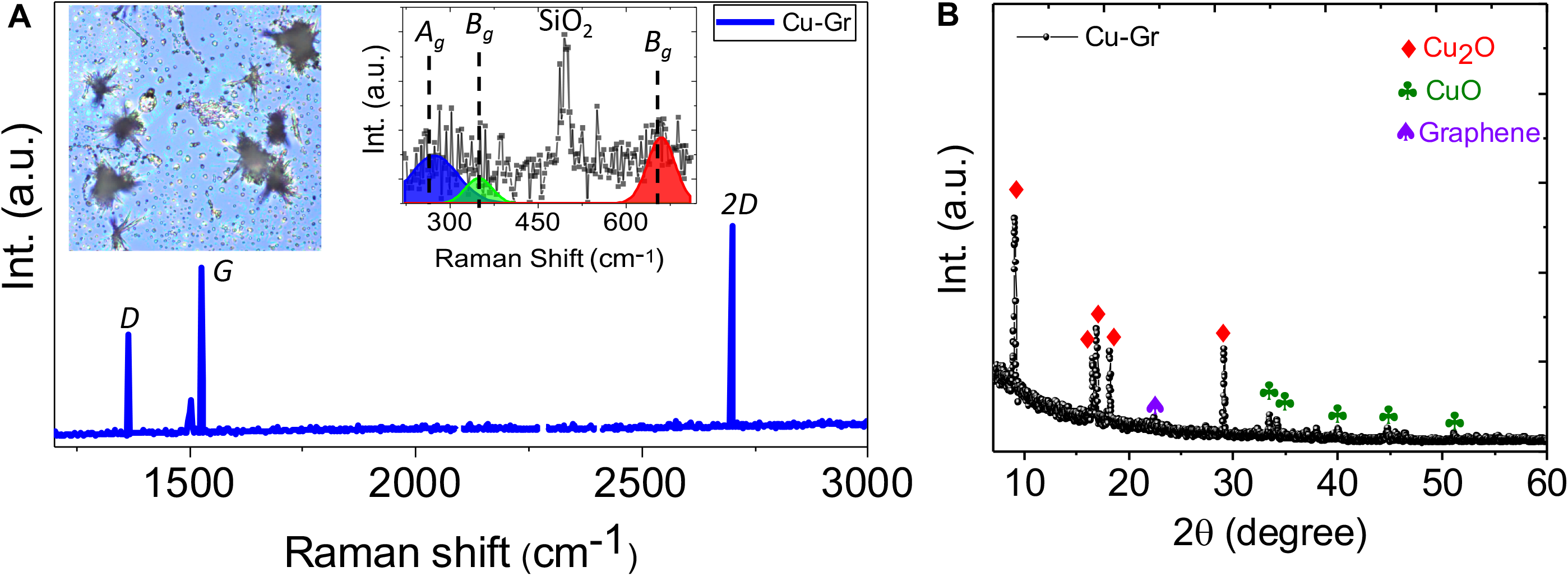
Material characterization of the Cu-Gr nanocomposite. (A) Raman spectrum of Cu-Gr nanocomposites. Inset shows optical microscopy image of the composite and Raman spectrum of Cu_x_O. (B) XRD spectrum of composites sample confirming the presence of CuO, Cu_2_O, and Graphene.

### Extensive characterization of antiviral property of Colloidal Cu-Gr nanocomposite

Followed by the material characterization, we have invested significant efforts for the characterization of the antiviral property of the Cu-Gr nanocomposites in its colloidal form. First, we have used the Nano-Luc reporter assay in order to identify the optimal time and concentration required for its antiviral activity. The Nano-Luc reporter assay was performed where the influenza A reporter virus was pretreated with the colloidal form of the Cu-Gr nanocomposite for various time periods before using them for infecting MDCK cells. As evidenced from Figure 5A, a sharp decrease (>50%) in the reporter activity was observed as a result of 30 minutes of preincubation with Cu-Gr composite, while longer times of preincubation showed only minor additional reduction. This data suggested that 30 minutes of preincubation with Cu-Gr composite can lead to more than tenfold reduction in input virus titer that ultimately results in ~50% decrease in reporter activity. Subsequently, we tried to identify the optimal concentration of the Cu-Gr composite required for its antiviral activity. Different concentrations of the Cu-Gr composite (50nM, 100nM, 500nM, 1μM, 2μM and 5μM, respectively) were used to treat the Nano-Luc influenza A reporter virus for 30 minutes followed by performing Nano-Luc reporter assay with the same. A precise dose dependent decrease in reporter activity and hence virus replication was observed as a result of prior treatment with Cu-Gr composite within the concentration range of 0.5mM to 5mM (Figure 5B). While higher concentration (10μM) further reduced reporter activity, it may also show cytotoxicity upon the cells, hence excluded from the subsequent experiment.

**Figure 5.**
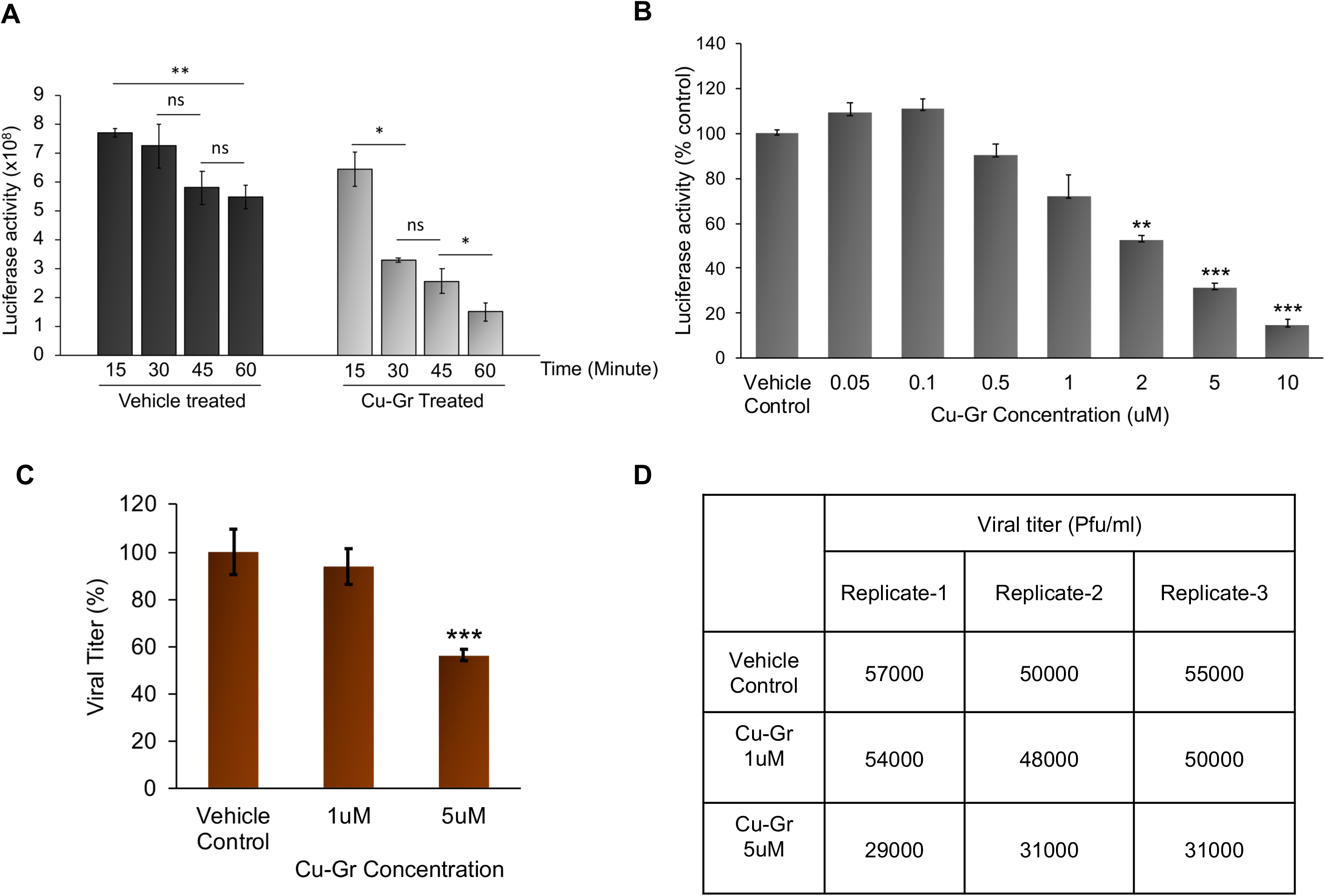
Prior incubation with Cu-Gr nanocomposite severely impacts the infectivity of influenza A virus. (A) A time kinetics experiment was performed by treating Influenza A reporter virus with 5uM colloidal suspensions of Cu-Gr for 15, 30, 45 and 60 min at RT and followed by infection of MDCK cells at MOI 0.1. Absolute luciferase activity values w.r.t viral gene expression for the vehicle and Cu-Gr treated sets are represented by black and grey bars respectively. (B) MDCK cells were infected with Influenza A reporter virus pretreated with vehicle or with different concentrations of the Cu-Gr composite (50nM, 100nM, 500nM, 1uM, 2uM and 5uM respectively). Viral gene expression was monitored using luciferase activity assay. Data were normalized to vehicle control sets for each nanoparticle. (C, D) A non-reporter A/H1N1/WSN/1933 influenza virus was either treated with Cu-Gr composite (1uM and 5uM) or with vehicle control prior to infecting MDCK cells. Plaque assay was performed to measure the titer of the progeny virus particles harvested at 8hpi. Percentage reduction in viral titers and actual PFUs for all experiments are represented. For all experiments, data are mean of n=3± standard deviation (*, P< 0.05; n=3 ± sd).

Next, we intended to test whether the reduction in the luciferase activity of the reporter virus, as a result of pretreatment with Cu-Gr nanocomposite, can also be correlated to the reduction of progeny virus titer. A non-reporter variant of the influenza A/H1N1/WNS/1933 virus was used for this purpose. Virus stock solutions were either treated with two different concentrations of Cu-Gr nanocomposites (1μM and 5μM) or with the vehicle control prior to infection on MDCK cells. Plaque assay was performed to measure the titer of the progeny virus particles harvested at 8 hours of post-infection. There is about 40% decrease in viral titer for the sets treated 5uM Cu-Gr solution with respect to the vehicle treated sets. Treatment with 1μM Cu-Gr nanocomposite shows non-significant decrease in viral titer. This data further substantiates the fact that treatment with 5μM Cu-Gr significantly reduces viral infectivity which results in a decrease in viral gene expression, replication and subsequent production of viral titer (Figure 5C). The plaque assay titer data are tabulated in Figure 5D.

### Prior treatment with Cu-Gr nanocomposite explicitly inhibits virus entry into the cells

At this point, we sought to examine the molecular mechanism by which Cu-Gr nanocomposite interferes with virus replication cycle. Metal nanoparticles have been shown to interfere with the integrity of the virus particles or the activity of the surface glycoproteins that may interfere with the entry of virus particles into the host cells (Sunada, Minoshima, and Hashimoto 2012) (Ting Du. 2018). Hence, to investigate the effect of pretreatment of Cu-Gr specifically upon virus entry step, we have performed an ‘entry assay’ (Figure 6A). The non-reporter variant of Influenza A WSN virus were either treated with Cu-Gr or with solvent and subsequently used to infect MDCK cells in a synchronized fashion. Post entry, cells were incubated with cycloheximide containing media for one hour to allow the import of the incoming viral ribonucleoprotein complexes (RNPs) into the nucleus. Subsequently, the incoming viral RNPs were stained with antibodies specific to viral Nucleoprotein (NP), which is the major component of the RNPs. As shown in the Figure 6 B, input viral RNPs are solely detected inside the nucleus of the infected cells irrespective of the treatment. However, prior exposure to the Cu-Gr composite resulted in a significant reduction in the number of NP positive cells. A quantitative analysis of 5 different fields with a total of 500 cells for both treated and untreated sets shows roughly 80% decrease in number of NP positive cells in the Cu-Gr treated set, with respect to the untreated one (Figure 6C). This data clearly suggests that exposure to Cu-Gr nanocomposite compromises the ability of the virus particles to enter into the host cell, possibly by impacting the structural integrity of the of the virion particles.

**Figure 6.**
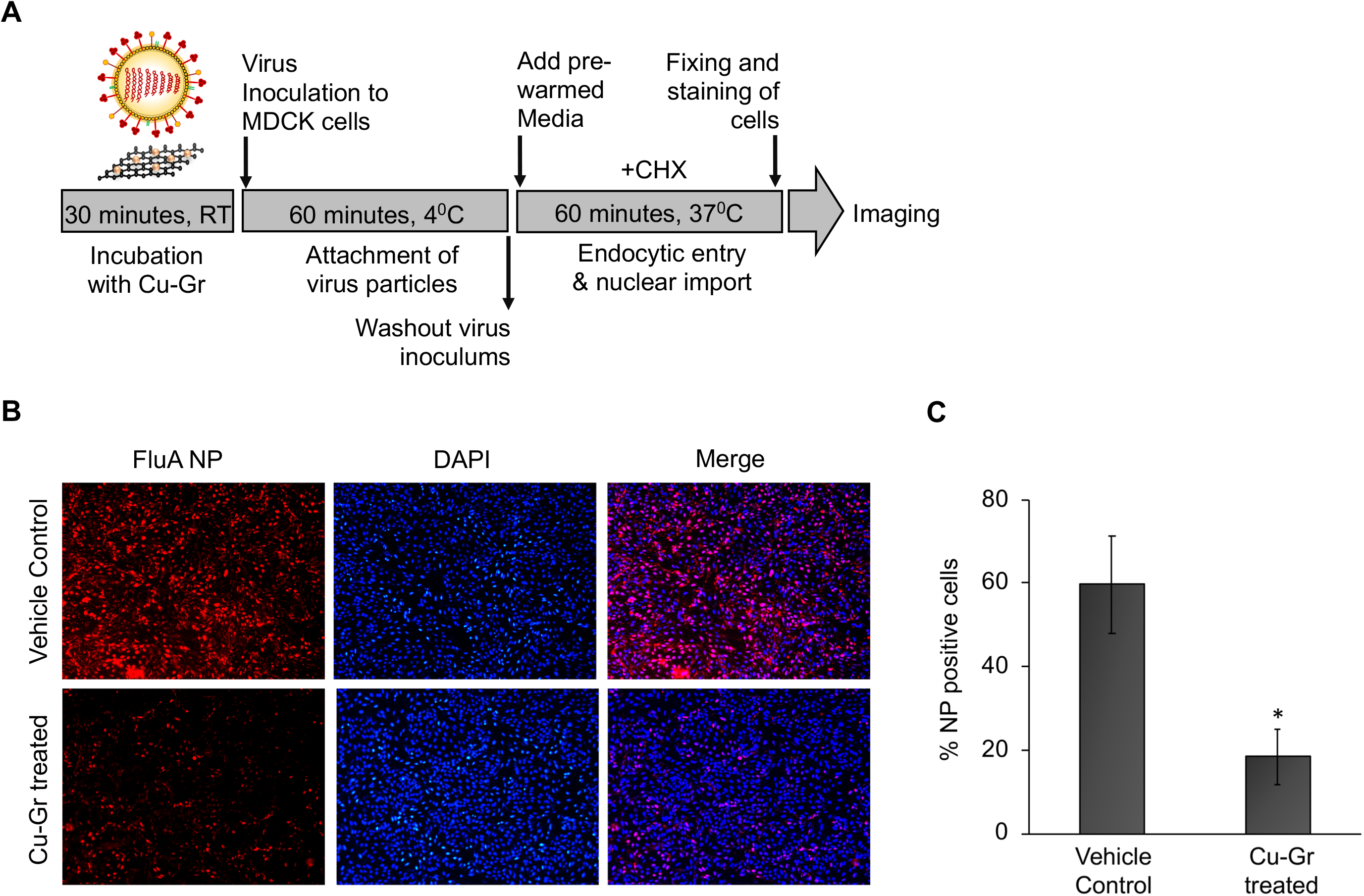
Cu-Gr interferes with the entry of Influenza A virus inside the cells. (A) Schematic depiction of Entry assay: MDCK cells were infected with influenza A virus either pretreated with 5uM Cu-Gr or with vehicle control. Synchronous infection was carried out by incubating the virus inoculum with the cell monolayer at 4°C for one hour, followed by adding the virus growth media (VGM) supplemented with cycloheximide and prewarmed at 37°C. Cells were incubated further for one hour before processing them for imaging (B) Intracellular localization of viral NPs in control and Cu-Gr treated sets were determined by staining with anti NP antibody and Alexa Fluor 555 (red). DAPI was used to stain the nucleus (Blue). (C) Percentage of NP positive cells in control and in treated sets were analyzed using image J software and depicted by bar diagram. For all experiments, data are mean of n=5± standard deviation (*, P< 0.05).

### Development of a Cu-Gr nanocomposite based transparent coating with strong antiviral activity

The ability of Cu-Gr nanocomposite to reduce the infectivity of influenza A virus prompted us to test its ability to inactivate influenza virus in the solid form. For this purpose, we have coated different wells of a 48 well plate with a series of coating solutions containing different concentrations of Cu-Gr composite (1μM, 5μM, 10 μM and 20 μM) and polyvinyl alcohol (PVA) (1mm, 5mm, and 10mm) as a capping agent. As a control, wells were coated with only PVA. This process generated a thin transparent film of Cu-Gr nanocomposite onto the surface of each well. To test the antiviral activity of these films, defined amounts of Nano-Luc influenza A reporter virus were inoculated in these coated wells and incubated for 30 minutes. Post treatment, infectivity of these virus inoculums were tested on MDCK cells using the Nano-Luc reporter assay as mentioned above. Prior exposure to the coating materials having various concentrations of Cu-Gr composite blended in different amounts of PVA resulted in differential effects upon viral replication (Figure 7). Films containing different concentrations of Cu-Gr composite, either in absence or in presence 1mM PVA barely showed any impact upon the infectivity of the virus. In contrast, an exact dose-dependent decrease in viral gene expression was observed as a result of prior treatment with the films containing increasing concentrations of Cu-Gr Composite in 5mM PVA. This antiviral activity was even more pronounced (70% decrease in Nano-Luc activity) for the films containing 1-5uM of Cu-Gr with 10mM of PVA. Together, this data shows 5uM of Cu-Gr composite capped with 10mM of PVA could be used to generate a transparent coating with high antiviral activity.

**Figure 7.**
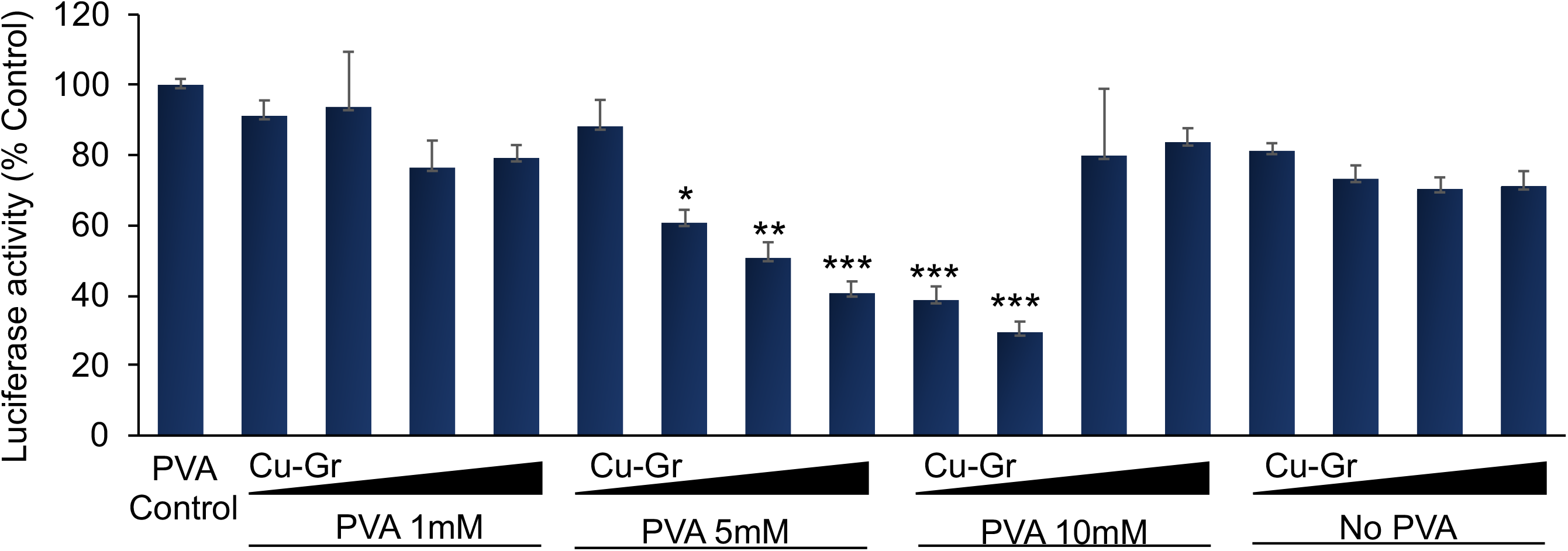
PVA based Cu-Gr nanocomposite coating shows strong antiviral activity. Different wells of a 48 well plate was coated with coating solutions having different concentrations of Cu-Gr composite (1uM, 5 uM, 10 uM and 20 uM) and polyvinyl alcohol (PVA) (1mM, 5mM and 10mM). Nano Luciferase-Influenza A virus (MOI 0.1) were inoculated in these wells and incubated for 30 minutes. Post treatment, MDCK cells were infected with the viral inoculum recovered from the coated wells and luciferase activity was determined at 8 hpi using Nano-Glo reporter assay (Promega). Luciferase activity of each set was plotted as a relative percentage of the vehicle treated set. Each data was represented in triplicate and each experiment was repeated three times. Data are presented as means ± standard deviations (SD) (n=3) (*, P< 0.05).

Encouraged by the results mentioned above, we have used dip-coating method used to coat a tempered glass with the Cu-Gr solution with optimum concentration. The glass unit was kept to soak the solution for 24 h and then air-dried naturally as shown in (Figure 8A). No formation of visibly aggregated spots or clogging was observed on the glass surface. A clean, transparent screen was obtained, and when fixed on a cell phone, there was no compromisation of light intensity or clarity of image on the display screen observed (Figure 8B). Optical transmittance spectra confirm the transparency of the coating on glass substrates (Figure 8C). As observed from the SEM images (Figure 8D), the Cu-O nanoparticles were uniformly embedded on top of the graphene layer. Elemental analysis of Cu-Gr compounds was conducted by color mapping and EDAX analysis. The data presented in the right panel of Figure 8D and Figure 8E confirmed the presence of C, O, and Cu elements in the composites sample and their uniform distribution in the sample. Figure 8F shows a schematic representation of the Cu_2_O and CuO nanoparticles embedded on the graphene sheets with PVA as a binding agent.

**Figure 8.**
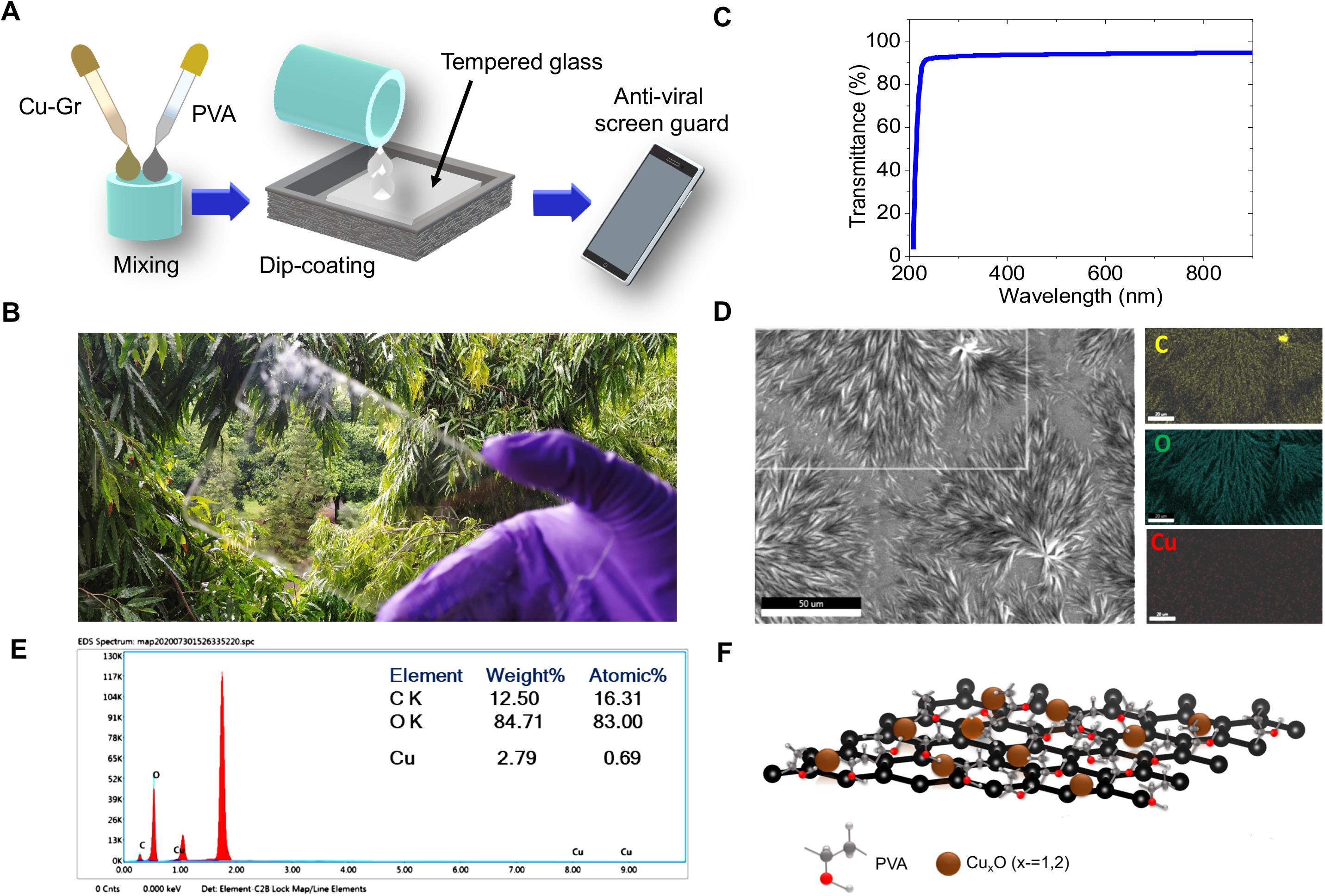
PVA based Cu-Gr nanocomposite can be used to generate a highly transparent antiviral coating of the cell phone screen. (A) Schematic diagram of the deep coating method of tempered glass unit with the PVA based Cu-Gr nanocomposite. (B) Transparency of dip coated tempered mobile screen. (C) Transmittance spectrum of the coated sample. (D) SEM image of Cu-Graphene composites sample. Right panel shows the color mapping of all elements. (E) EDX spectra of composites sample, confirm the presence of Cu, C and O atoms. (F) Schematic representation of the composite structure.

## Discussions

RNA viruses constitute one of the broader families of human pathogens, including influenza, Nipah, Ebola, SARS or MERS-Coronaviruses. Irrespective of their specific differences in virus replication cycle, all of these viruses share broader structural similarities. The viral genomic RNA remains enwrapped with single or multiple viral proteins and remains enclosed within the lipid bilayer envelope embedded with viral spike proteins. The integrity of the lipid envelope and the functionality of the spike proteins are not only crucial for the protection of the viral genomic materials in the outer environment but also indispensable for conducting the first two steps of the virus life cycle that are “attachment” and “entry” (Jane Flint, Vincent R 2015). In this study, we have identified Cu-Gr nanocomposite as a potential antiviral agent that can interfere with these two steps of the influenza A virus life cycle, possibly through compromising the structural integrity of the virion particles.

We, for the first time, have used a highly sensitive Nano-Luc reporter assay to perform an unbiased screening of a series of nanoparticles and their composite materials in order to identify a coating substance with strong antiviral activity. Our data identified Cu-Gr nanocomposite as the most potential antiviral agent. Interestingly, Cu nanoparticles or graphene individually showed minimal or no antiviral activity in our assay while a hybrid between these two showed at least ten-fold decrease in effective viral titer. This might be due to the property of the composite substances, which is not just a hybrid between two materials in their original state, but rather a combination of the modified version of the materials (Ramakrishnan et al. 2015). This can also be substantiated from the previous studies showing metal-ion based composites shows better activity compared to that of metal nanoparticles itself (Minoshima et al. 2016; Perdikaki et al. 2018; Sunada, Minoshima, and Hashimoto 2012). Studies also reveal that the antiviral activity mainly depends on the presence of ions generated from the surface of nanoparticles (Ma et al. 2011) (Shen et al. 2010). Our XRD data clearly indicates the presence of various copper oxide species, Cu_2_O and CuO, embedded in the graphene sheets which may serve as the basis for the antiviral activity of the composite material. The presence of reactive oxygen species in Cu_x_O-graphene sheets may provide ions, which may interfere with the structural integrity of the lipid bilayer membrane or the surface antigens of the virion particles. This interference should compromise the ability of the virus particles to interact with the cell surface receptor essential for the attachment and subsequent entry into the host cell. Our entry assay supports this hypothesis, as prior treatment with Cu-Gr composite resulted in significant reduction in the percentage of infection positive cells with respect to the vehicle treated set.

Finally, we have used PVA as a capping agent and identified optimum concentrations of Cu-Gr and PVA to develop a thin transparent coating with intense antiviral activity. While we have used dip-coating method to coat a tempered glass with high visibility, other forms of coatings like doctor’s blade technique, spin coating, and spray coating can also be implemented. Due to the high transmission efficiency, such coating material could be implemented on a wide variety of surfaces, which could radically decrease the stability of the virus particles in the outer environment and hence reduce the transmission rate drastically. Needless to mention that such generic antiviral strategy can significantly reduce the overall burden of seasonal respiratory infection-related epidemics or occasional pandemics caused either by various influenza or coronaviruses.

## Materials and Methods

### Chemicals

Copper sulfate (CuSO_4_ > 99%), Silver nitrate (AgNO_3_ > 99%), Sodium borohydride (NaBH_4_ > 98%), Poly vinyl alcohol (>99%), Sodium hydroxide pellets (NaOH > 97%) and graphite powder (>98%) were purchased from Sigma-Aldrich.

### Cells, Viruses and Antibody

Madin Darby Canine Kidney (MDCK) (CCL-34) cells were maintained in Dulbecco’s modified Eagle’s medium (DMEM) supplemented with 5% FBS at 37°C and 5% CO_2_ along with penicillin and streptomycin antibiotics (Gibco).

Influenza A virus strains, A/WSN/1933 (H1N1), WSN stably encoding PB2 with a C-terminal FLAG tag (WSN-PB2-FLAG) (Dos Santos Afonso et al. 2005) or PA-2A-Swap-Nluc (PASTN) reporter virus based on the strain A/WSN/33 (H1N1) were used for infecting the cells (Tran et al. 2013). Antibody used includes anti-NP (H16-L10-4R5) (Yewdell et al., 1981).

### Synthesis of materials

Initially, the graphene dispersion was prepared using a liquid exfoliation of the graphite powder in Deionized (DI) water (20 mg/300 mL) using an ultrasonic probe sonicator. Probe sonication of frequency 30 Hz was used in pulses for 2 hours. Silver (Ag) and Copper (Cu) stock solutions were prepared using 4mM AgNO_3_ (68 mg/100 mL) and 4 mM CuSO_4_ (63.8 mg/100 mL) in DI water medium respectively. Graphene solution (45 mL) was mixed with 15 mL of Cu and Ag stock solution separately by maintaining the pH=12 adjusted through NaOH. 20 mL of 4 mM NaBH_4_ (30 mg/200 mL) solution, a strong reducing agent is added drop wise with the Cu-graphene (Cu-Gr), Ag-graphene (Ag-Gr) and Cu-Ag-graphene (Cu-Ag-Gr) mixtures separately and stirred continuously at 40°C.

Furthermore, four different concentrations (1μM, 5μM, 10μM and 20μM) of Cu functionalized graphene samples were synthesized for biological process optimization. Three different concentrations (1mM, 5mM and10mM) of poly vinyl alcohol have been capped as a coating media for the Cu-graphene samples.

### Material characterizations

Different phases of the synthesized sample (Cu functionalized graphene) were obtained from the X-ray diffraction (XRD) peaks by using Bruker D8 Advance X-ray diffractometer within a scan range of 2Theta (2θ) values 7 and 90° with Cu-Kα source, maintaining the scan rate of 1° min^−1^. Absorption spectra of Cu-graphene synthesized sample were recorded by a BioTek UV-vis spectrophotometer Epoch 2 microplate within 200 nm to 800 nm wavelength. Raman shifts were measured by using WiTec –alpha 300R confocal microscope at excitation of 532 nm in the wavenumber range of 200 cm^−1^ to 3000 cm^−1^. SEM images were obtained through Zeiss-Merlin EVO 60 scanning electron microscope with Oxford EDS detector.

### MTT assay

MDCK cells were seeded in 96 well plates at a density of 15000 cells per well. The cells were treated with Silver (Ag), Graphene (Gr), Copper (Cu) nanoparticles as well as Ag-Gr, Cu-Gr and Ag-Cu-Gr nanocomposites respectively for 24 hours (h) at 37°C in 5% CO_2_. Cellular cytotoxicity assay was performed according to the manufacturer’s protocol. In brief, after 24 h of treatment with the nanoparticle, 100μl of MTT reagent (5 mg/ml, SRL) in PBS was added to the cells and incubated for 3 h at 37°C. The medium was removed carefully without disturbing the formazan crystals and 100μl of DMSO (Sigma) was added to dissolve the insoluble purple formazan crystals. The absorbance of the suspension was measured at 595 nm using Epoch 2 Microplate Reader (BioTek instruments). The percentages of metabolically active cells were compared with the percentage of control cells of the same culture plate as a proxy for cell viability. Cellular cytotoxicity was determined in triplicate and each experiment was repeated three times independently.

### Nano-Luc reporter assay

The Nano-Luc reporter assay was used to determine the luciferase activity as previously mentioned by Tran et al., 2013 (Tran et al. 2013). MDCK cells seeded in a 96 well plate were infected in triplicate with the Nano Luciferase influenza A reporter virus, PASTN. Accordingly, the virus was preincubated with a particular concentration of the nanoparticle/nanocomposite for 30 min or mentioned otherwise. The vehicle control or the nanoparticle/ nanocomposite treated virus was used to infect MDCK cells. The infected cells were harvested at 8hpi and the viral NLuc activity was measured by using Nano-Glo® Luciferase Assay System according to the manufacturer’s instructions (Promega) and the luminescence was detected by using a Luminometer (Glomax 20/20, Promega).

### Plaque assay

A non-reporter variant of the influenza A/H1N1/WSN/1933 virus stock solutions MOI 0.1 was either preincubated with vehicle control or Cu-Gr nanocomposite (1uM and 5uM) followed by infection in MDCK cells. At 8hpi, the viral supernatant was collected and used to reinfect MDCK cells and plaque assay was performed to determine the progeny viral titer followed by Matrosovich M, 2006 (Matrosovich et al. 2006). Accordingly, after 1 hr of infection with the virus, cells were overlaid with a media containing a mixture of 2X DMEM and 2.4% avicel (1:1 ratio). After 62hrs, the overlay was discarded and the cells were fixed with 70% ethanol followed by staining with 2% Crystal violet. Plaques were counted and plaque forming unit (PFU/ml) was calculated to measure the progeny viral titer.

### Viral entry assay

MDCK cells grown on coverslips were infected with vehicle control or 5uM Cu-Gr treated virus at a MOI of 5. Viral entry assay was performed according to the protocol followed by Mondal et al., 2017 (Mondal et al. 2017). Virion binding was performed at 4°C for 1hr in presence of 1mM of Cycloheximide, CHX (Sigma). The viral inoculum was washed off with cold PBS to remove unbound virus particles. Thereafter, the prewarmed virus growth media (VGM, containing DMEM, 0.2% bovine serum albumin (BSA), 25 mM HEPES buffer, and 0.5 mg/ml TPCK-trypsin) supplemented with 1mM of CHX was added to the cells and synchronous infection was initiated by shifting cells to 37°C. At 1 hpi, cells were fixed with 3% formaldehyde and permeabilized with 0.1M Glycine/0.1% Triton-X 100 in PBS for 20 min at room temperature. Blocking was performed at 4°C with 3% BSA overnight. NP was detected with anti-NP antibody and Alexa Fluor 555-conjugated donkey anti-mouse IgG antibody (Invitrogen). DAPI (Sigma) was used to stain the nucleus. Cells were imaged using Fluorescent microscope (Leica Microsystems) and image analysis was performed with ImageJ software.

### Statistics

Each data is a representative of at least three independent experiments, each experiment was performed in triplicate. Graphs are performed in Microsoft Excel and represented as mean standard deviations (n=3). Results were compared by performing two-tailed Student’s t test. Significance is defined as p<0.05 and statistical significance is indicated with an asterisk (*). The *p value < 0.05, **p value < 0.01 **p value and ***p < 0.001 were considered statistically significant.

## Acknowledgement

We sincerely thank Prof. Andrew Mehle, University of Wisconsin Madison, for providing the Nano-Luc influenza A reporter viruses. This work was primarily supported by Sponsored Research and Industrial Consultancy (SRIC) IIT Kharagpur. AM would like to thank DBT for Ramalingaswami re-entry fellowship, SERB for Early Career Research Award and MHRD for the “Scheme for Transformational and Advanced Research in Science” for additional financial support.

